# Manganese/iron-supported sulfate-dependent anaerobic oxidation of methane by archaea in lake sediments

**DOI:** 10.1101/636621

**Authors:** Guangyi Su, Jakob Zopfi, Haoyi Yao, Lea Steinle, Helge Niemann, Moritz F. Lehmann

## Abstract

Anaerobic oxidation of methane (AOM) by methanotrophic archaea is an important sink of this greenhouse gas in marine sediments. However, evidence for AOM in freshwater habitats is rare, and little is known about the pathways, electron acceptors and microbes involved. Here, we show that AOM occurs in anoxic sediments of a lake in southern Switzerland (Lake Cadagno). Combined AOM-rate and 16S rRNA gene-sequencing data suggest that *Candidatus* Methanoperedens archaea are responsible for the observed methane oxidation. Members of the Methanoperedenaceae family were previously reported to conduct nitrate- or iron/manganese-dependent AOM. However, we demonstrate for the first time that the methanotrophic archaea do not necessarily rely upon these oxidants as terminal electron acceptors directly, but mainly perform canonical sulfate-dependent AOM, which under sulfate-starved conditions can be supported by metal (Mn, Fe) oxides through oxidation of reduced sulfur species to sulfate. The correspondence of high abundances of Desulfobulbaceae and *Candidatus* Methanoperedens at the same sediment depth confirm the interdependence of anaerobic methane-oxidizing archaea and sulfate-reducing bacteria. The relatively high abundance and widespread distribution of *Candidatus* Methanoperedens in lake sediments highlight their potentially important role in mitigating methane emissions from terrestrial freshwater environments to the atmosphere, analogous to ANME-1, -2 and -3 in marine settings.

## Introduction

A major fraction of the methane (CH_4_) produced in lakes is oxidized by methanotrophic bacteria right at the redox transition zone within sediments or in the water column (Bastviken et al. 2002; Blees et al. 2014). Yet, more recent reports indicate that methane in lakes is also oxidized in the absence of oxygen (Schubert et al. 2011; Sivan et al. 2011; Deutzmann et al. 2014; Martinez-Cruz et al. 2018). Information on the controls on lacustrine methane oxidation in general, and on the alternative electron acceptors involved in anaerobic oxidation of methane (AOM) in particular, is important for understanding the regulation of the biological methane filter in lakes (Sivan et al. 2011; Norði et al. 2013; Deutzmann et al. 2014; Weber et al. 2017).

AOM coupled to sulfate reduction has been recognized as the most important sink in marine environments, where sulfate concentrations are high (Reeburgh 2007; Knittel and Boetius 2009). This microbial process is primarily mediated by anaerobic methanotrophic archaea (ANME), and sulfate-reducing bacteria (SRB) (Boetius et al. 2000; Orphan et al. 2002; Niemann et al. 2006; Milucka et al. 2012; Wegener et al. 2015). AOM, putatively coupled to sulfate reduction, has also been observed recently in freshwater ecosystems, for example wetlands (Segarra et al. 2015), iron-rich lake sediments (Norði et al. 2013; Weber et al. 2017), and ditch sediments (Timmers et al. 2016). However, in most lacustrine environments (e.g., in anoxic lake sediments), sulfate-dependent AOM is likely limited by relatively low sulfate concentrations.

Possible alternative terminal electron acceptors for AOM include nitrate and nitrite. Indeed, both nitrate- and nitrite-dependent AOM have recently been documented in laboratory enrichment cultures or freshwater systems (Raghoebarsing et al. 2006; Ettwig et al. 2010; Haroon et al. 2013; Deutzmann et al. 2014). Moreover, AOM coupled to the reduction of metal oxides (i.e., ferrihydrite and birnessite) has been demonstrated in anoxic marine sediments (Beal et al. 2009), a freshwater enrichment culture (Ettwig et al. 2016), and in lake sediments (Sivan et al. 2011; Norði et al. 2013). Despite the potential for iron- and manganese-coupled AOM as major methane sink in many Fe/Mn-rich sedimentary environments, the electron transport mechanisms that couple AOM with metal oxides (as well as sulfate) are still not fully understood (Milucka et al. 2012; McGlynn et al. 2015; Wegener et al. 2015). Moreover, it remains unclear whether microorganisms in environments where both metal oxides and sulfate are present (Egger et al. 2015; Weber et al. 2017) can independently mediate AOM using iron or manganese oxides (i.e., Fe(III)/Mn(IV)) as the terminal electron acceptors (Ettwig et al. 2016; Cai et al. 2018), or whether canonical sulfate-dependent AOM is indirectly stimulated by metal oxides that drive sulfide/sulfur oxidation via a cryptic sulfur cycle (Holmkvist et al. 2011a; Hansel et al. 2015).

Despite growing evidence that anaerobic methanotrophs play an important role in removing methane from lake ecosystems and reducing fluxes to the atmosphere, our knowledge about the microorganisms that perform AOM, particular within lake sediments, is still rudimentary. So far, only a few studies have identified freshwater AOM-mediating microorganisms (Ettwig et al. 2010, 2016; Schubert et al. 2011; Haroon et al. 2013; Weber et al. 2017; Graf et al. 2018; Versantvoort et al. 2018). In freshwater lake sediments, methanotrophic bacteria related to *Candidatus* Methylomirabilis oxyfera have been reported to perform methane oxidation coupled to denitrification (Deutzmann et al. 2014), and archaea within the phylum Euryarchaeota possibly carried out sulfate- and/or iron-dependent AOM (Schubert et al. 2011; Weber et al. 2017). Although anaerobic methanotrophic archaea (e.g., ANME-2a) are potentially versatile with regards to the mode of AOM (Wang et al. 2014), the biogeochemical controls on possible metabolic adaptions in lacustrine environments are still poorly understood.

In the present study, we investigated methane oxidation in the anoxic sediments of Lake Cadagno. Using a complementary approach combining radio-tracer techniques (^14^CH_4_) for rate determination, incubation experiments with ^13^C-labeled methane and different electron acceptors and stable isotope probing (SIP) of lipid biomarkers, as well as 16S rRNA gene sequencing, we aimed at 1) revealing the microbial processes and mechanisms involved in AOM with particular focus on the potential role of metal oxides in stimulating sulfate-dependent AOM, and 2) identifying the freshwater AOM-mediating microorganisms responsible for methane oxidation within the sediments. We demonstrate that methane oxidation is primarily coupled to sulfate reduction (even at sediment depths where sulfate is depleted), yet AOM cannot be attributed to the typical ANME lineages that were found to perform AOM with sulfate (Knittel and Boetius 2009), but is mediated by thus far uncultured archaea (Schubert et al. 2011), related to *Candidatus* Methanoperedens (formerly named ANME-2d or AAA).

## Materials and Methods

### Study Site

Lake Cadagno is an alpine meromictic lake located in the southern Alps of Switzerland (46°33′N, 8°43′E). The permanent chemocline in the water column of this lake separates the oxic mixolimnion from the anaerobic and sulfidic monimolimnion. Due to water infiltration from high-ionic strength subaquatic springs, Lake Cadagno displays relatively high concentrations of sulfate (>1 mmol/L).

### Sampling

A total of six undisturbed sediment cores (inner diameter 62 mm) were recovered with a gravity corer from the deepest site (21 m water depth) in Lake Cadagno in October 2016, and subsampled in the home laboratory for geochemical analyses and AOM rate measurements. Samples for dissolved methane concentrations were collected onsite with cut-off syringes through holes in one of the core tubes that were covered with tape during coring. 3 mL of sediment samples were fixed with 7.0 mL 10% NaOH in 20 mL glass vials, which were then immediately sealed with thick butyl rubber stoppers (Niemann et al. 2015). A second sediment core was sacrificed for the quantification of sulfur species, dissolved and particulate iron/manganese, as well as for DNA extraction. The sediment core was sectioned into 1 or 2 cm segments, and DNA samples were collected and stored frozen at −20 °C until further processing. Pore water was extracted by centrifuging the sediment samples under anoxic condition, and filtering the supernatant through 0.45 µm filters. Porewater samples (200 µL) for sulfide concentration measurements were fixed with Zn acetate (5% w/v) immediately after filtration. For the analysis of dissolved iron and manganese concentrations, 1 mL of the filtered sample was amended with 200 µL 6 M HCl. For the analysis of dissolved inorganic carbon (DIC), sulfate and nitrogen species concentrations, the remaining samples were stored at 4 °C, respectively.

### Porewater and sediment geochemical analyses

Methane concentrations in the headspace of NaOH-fixed samples were measured using a gas chromatograph (GC, Agilent 6890N) with a flame ionization detector, and helium as a carrier gas. The C isotopic composition (^13^C/^12^C) of methane from the headspace was determined using a pre-concentration unit (Precon, Finnigan) connected to an isotope ratio mass spectrometer (IRMS; Delta XL, Finnigan). Stable C-isotope values are reported in the conventional δ notation (in ‰) relative to the Vienna Pee Dee Belemnite standard (V-PDB). δ^13^CH_4_ values have an analytical error of ± 1%. A total carbon analyzer (Shimadzu, Corp., Kyoto, Japan) was used to quantify dissolved inorganic carbon (DIC) concentrations in the porewater. Hereby, DIC was quantified as the difference between the total dissolved carbon concentration and the dissolved organic carbon concentration, which was analyzed separately after acidification of the sample with HCl. Porewater concentrations of ammonium, sulfate and nitrate were analyzed by ion chromatography (Metrohm, Switzerland). Sulfide concentrations were determined spectrophotometrically using the Cline method (Cline 1969). Dissolved iron (Fe(II)), manganese (Mn(II)), and total manganese concentrations were measured by inductively coupled plasma optical emission spectrometry (ICP-OES). Reactive Fe(III) (FeOx) in the solid phase was extracted with 0.5 M HCl and then reduced to Fe(II) with 1.5 M hydroxylamine. Concentrations of Fe(II) were then determined photometrically using the ferrozine assay (Stookey 1970). Particulate reactive iron was calculated from the difference between the total Fe(II) concentrations after reduction, and the dissolved Fe(II) in the filtered sample.

### Flux calculations

Diffusive fluxes *J* (in mmol/m^2^/s) of methane and sulfate in the sediment porewater were calculated according to Fick’s first law of diffusion, assuming steady-state conditions (Eq. 1):

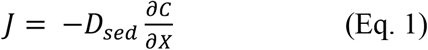

where *D*_*sed*_ is the molecular diffusion coefficient D_0_ (in m^2^/s) for methane and sulfate, respectively, corrected for sediment porosity (0.93) and the corresponding tortuosity. 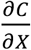is the solute concentration gradient (in mmol/m^4^), which was estimated based on the linear portions of concentration profiles within the investigated depth intervals. Molecular diffusion coefficients D_0_ were adopted from Boudreau (1997), under consideration of the in situ temperature in Lake Cadagno sediments (9.48×10^−11^ and 5.08×10^−10^ m^2^/s for methane and sulfate, respectively).

### AOM Rate Measurements

A radiotracer ^14^CH_4_ technique (Iversen and Jørgensen 1985) was chosen to obtain depth-specific ex-situ AOM rate profiles in the sediments. ^14^CH_4_ radiotracer injection was applied directly to the whole core through pre-drilled side-holes at a depth interval of 2 cm. Subsequent incubations were performed at in situ temperature (4 °C) in the dark. At the end of the incubation, biological activity was stopped by adding aqueous NaOH (5% w:w). ^14^CH_4_ activity was measured in the residual methane (as CO_2_ after combustion), the CO_2_ produced by AOM, and the remaining biomass via liquid scintillation counting (Blees et al. 2014; Steinle et al. 2016). AOM first order rate constants (*k*) were calculated according to Eq. 2.

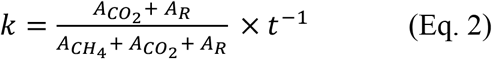

where 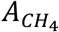, 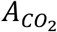 *and A*_*R*_ represent the radioactivity of methane, CO_2_, and the remaining radioactivity. *t* represents the incubation time. Methane oxidation rates (MOR) were then calculated using the value for k and the methane concentration at the start of the incubation (Eq. 3).

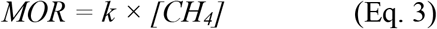

### ^13^CH_4_ Incubation Experiments

Sediments from three sediment zones: 14-19 cm (where maximum AOM rates were observed; Fig. 1A), 19-24 cm and 24-29 cm (where sulfate concentrations were below the detection limit) from four replicate cores were combined, each section with approximately 600 cm^3^ of fresh sediment, and mixed with 2.5 L anoxic artificial mineral medium (Ettwig et al. 2009) to yield homogenized mixtures for the slurry incubation experiments (see Table S1 for components of medium). 240 mL serum vials were filled with ~ 200 mL of the homogeneous sediment slurry. The slurries were degassed with He to remove any traces of O_2_ and background methane. All slurries were pre-incubated under anoxic condition for at least two weeks to allow the microbial community to recover from any potential perturbation during sample handling, and the supernatant was replaced with anoxic sulfate-free water (this step was repeated until sulfate concentrations were below detection limit) (Segarra et al. 2013). In a first set of experiments, we prepared incubations with sediments only from 14-19 cm and from 19-24 cm. All bottles except for the controls were amended either with nitrate, sulfate, amorphous manganese or iron oxide, with final concentrations of 4.8 mM, 2.4 mM, 10 mM and 10 mM, respectively (Table 1). Control experiments include live controls (slurries without additional electron acceptors), killed controls (autoclaved unamended slurries), and incubations with sulfate-reduction inhibition (with 20 mM molybdate). In a second set of experiments, we split the slurries from the first set of incubation experiments (Cadagno sediments 19-24 cm) at the end of incubation period (after 96 days), re-amended them with MnO_2_ or sulfate, and in addition added 20 mM molybdate to some of them, respectively. Similarly, we divided the live control and added 0.5 mM nitrate to one of the splits. All incubation bottles were supplemented with 20 mL pure ^13^CH_4_ (99.8 atom %, Campro Scientific) to the He headspace, and were incubated in the dark in an anoxic chamber with N_2_ atmosphere at 25 °C. At different time points (0, 4, 8, 16, 32, 48, 64, 96 days), the sample was homogenized and 5 mL of the supernatant was collected and filtered with a 0.45 µm membrane filter for subsequent sulfate, dissolved iron/manganese, DIC concentration and stable carbon isotope ratio analyses. To determine the carbon isotope composition of DIC, 1 mL of water sample was transferred into a 12 mL He-purged exetainer (Labco Ltd) containing 200 µL zinc chloride (50 % w/v), and then acidified with ~100 µL concentrated H_3_PO_4_. The liberated CO_2_ was analyzed in the headspace using a purge-and-trap purification system (Gas Bench) coupled to a GC-IRMS (Thermo Scientific, Delta V Advantage). Absolute ^13^C-DIC abundances were determined from the DIC concentrations and the ^13^C/^12^C ratios converted from δ^13^C-DIC values (Oswald et al. 2015). The temporal change in ^13^C-DIC with incubation time was then used to calculate slurry-incubation-based potential methane oxidation rates, and to compare rates between the different treatments (Table 1).

**Figure 1.**
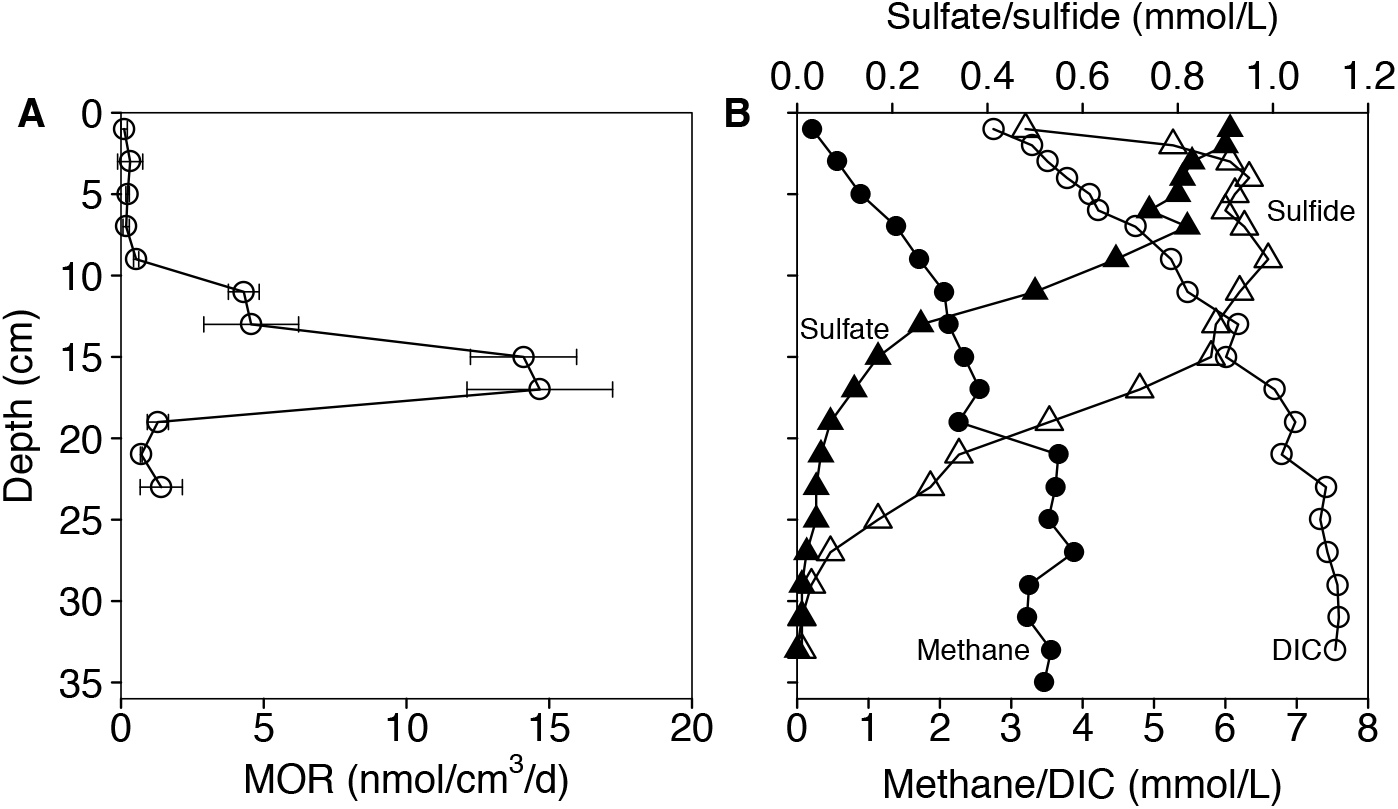
(A) Depth-specific in situ AOM rates determined by radioisotope-based approaches using whole-core incubations, (B) profiles of dissolved methane, DIC, sulfate and sulfide concentrations in the sediments of Lake Cadagno.

**Table 1.** ^13^CH_4_ incubation experiments with slurries from different sediment depths in Lake Cadagno.

| Cadagno sediment slurries |  | Treatment |  |
| --- | --- | --- | --- |
| Sampling interval* |  |  |  |
| 14-19 cm<br>and<br>19-24 cm<br>(first set of experiments) |  | Killed control | Autoclaved |
|  |  | Live control | - |
|  |  | FeOx<sup>†</sup> | +10 mM |
|  |  | MnO<sub>2</sub><sup>†</sup> | +10 mM |
|  |  | Sulfate | +2.4 mM |
|  |  | Nitrate | +4.8 mM |
|  |  | Molybdate | +20 mM |
| 24-29 cm<br>(second set of experiments) |  | Killed control | Autoclaved |
|  |  | MnO<sub>2</sub> | +10 mM |
|  |  | MnO<sub>2</sub> + Molybdate | +10 mM + 20 mM |
|  |  | Sulfate | +4.8 mM |
|  |  | Sulfate + Molybdate | +4.8 mM + 20 mM |
| 19-24 cm<br>(second set of experiments) |  | Live control | - |
|  |  | Nitrate | +0.5 mM |
|  |  | MnO<sub>2</sub> | +10 mM |
|  |  | MnO<sub>2</sub> + Molybdate | +10 mM + 20 mM |
|  |  | Sulfate | +4.8 mM |
|  |  | Sulfate + Molybdate | +4.8 mM + 20 mM |

### Microbial Lipid Extraction and Sample Analysis

Lipids were extracted from incubation slurries and further treated according to previously described methods (Elvert et al. 2003; Niemann et al. 2005). Briefly, total lipid extracts (TLEs) were obtained by ultrasonication of the slurry samples in four steps with solvent mixtures of decreasing polarity: (1) dichloromethane (DCM):methanol (MeOH) 1:2; (2) DCM:MeOH 2:1; (3) and (4) DCM. TLEs were then saponified with methanolic KOH-solution (12%) at 80 °C for 3 h. After extracting the neutral fraction, fatty acids (FAs) were methylated using BF_3_ in methanol, and analyzed later as FA methyl esters. The double-bond positions of monounsaturated fatty acids were determined by analyzing their dimethyl disulfide (DMDS) adducts (Nichols et al. 1986; Moss and Lambert-fair 1989). Neutral compounds were further separated into hydrocarbon, ketone and alcohol fractions using silica glass cartridges, followed by derivatization of alcohol fractions into trimethylsilyl ethers prior to analysis. Individual lipid compounds were quantified and identified by gas chromatography with flame ionization detection (GC-FID) and GC-mass spectrometry (GC-MS, Thermo Scientific DSQ II Dual Stage Quadrupole), respectively. Compound-specific stable carbon isotope ratios were determined using a GC-isotope ratio mass spectrometer (GC-Isolink Delta V Advantage, Thermo Scientific). Both concentrations and δ ^13^C values of lipids were corrected for the introduction of carbon atoms during derivatization. Accuracy and reproducibility of lipid concentrations and δ^13^C were monitored by repeated analysis of internal standards (n-C19:0-FA, n-C19:0-OC, α-Cholestane and n-C36:0). Reported δ ^13^C values have an analytical error of ± 1‰.

### DNA extraction, PCR amplification, Illumina sequencing and data analysis

DNA was extracted from both, samples of Lake Cadagno core sediments as well as from slurry sediments at the end of incubations, using a FastDNA SPIN Kit (MP Biomedicals) following the manufacturer’s instructions. A two-step PCR approach was applied in order to prepare the library (cf. Illumina support document 16S Metagenomic Sequencing Library Preparation (15044223 B)) for sequencing at the Genomics Facility Basel (https://www.bsse.ethz.ch/genomicsbasel). Briefly, a first PCR (25 cycles) was performed using universal primers 515F-Y (5′-GTGYCAGCMGCCGCGGTAA) and 926R (5′-CCGYCAATTYMTTTRAGTTT-3’) targeting the V4 and V5 regions of the 16S rRNA gene (Parada et al. 2016). Sample indices and Illumina adaptors were added in a second PCR of eight cycles. Purified indexed amplicons were finally pooled at equimolar concentration into one library and sequenced on an Illumina MiSeq platform using the 2×300 bp paired-end protocol (V3 kit). After sequencing, quality of the raw reads was checked using FastQC (v 0.11.8) (Andrews 2018). FLASH (Magoč and Salzberg 2011) was used to merge forward and reverse reads into amplicons of about 374 bp length, allowing a minimum overlap of 15 nucleotides and a mismatch density of 0.25. Quality filtering (min Q20, no Ns allowed) was carried out using PRINSEQ (Schmieder and Edwards 2011). Classical OTU (operational taxonomic unit) clustering with a 97 % cutoff was performed using the UPARSE-OTU algorithm in USEARCH v10.0.240 (Edgar 2013). Amplicon sequence variants were determined by denoising using the UNOISE algorithm (unoise3 command) and are herein referred to as ZOTU (zero-radius OTU). Taxonomic assignment was done using SINTAX v10.0.240_i86linux64 (Edgar 2016) and the SILVA 16S rRNA reference database v128 (Quast et al. 2013). Subsequent data analyses were carried out with Phyloseq (McMurdie and Holmes 2013) in the R environment (R Core Team, 2014) (http://www.r-project.org/).

### Data availability

Raw reads are deposited at the NCBI Sequence Read Archive (SRA) in the BioProject PRJNA497531, and can be accessed under the accession numbers SRR8130729-SRR8130745. Additionally, amplified sequence variants of *Candidatus* Methanoperedens and Desulfobulbaceae (ZOTU307) used to construct phylogenetic trees have been deposited in the GenBank, under the accession numbers MK087688-MK087694.

## Results and Discussion

### Anaerobic oxidation of methane in Lake Cadagno sediments

Methane oxidation has been investigated in Lake Cadagno sediments previously. Schubert et al. (2011) observed a strong ^13^C enrichment within the porewater methane pool close to the sediment-water interface, and attributed the elevated δ^13^CH_4_ values to the high C isotope fractionation associated with AOM. The apparent restriction of AOM hotspots to the uppermost 2-4 centimeters of the sediment column, where sulfate concentrations were up to 2 mM, led Schubert et al. (20111) to the conclusion that methane was most likely oxidized with sulfate as electron acceptor, and that AOM was constrained by the availability of sulfate in the sediment porewater. Here, we confirm the biogeochemical evidence for AOM in the Lake Cadagno sediments by providing, for the first time, downcore AOM rate measurements for these lake sediments. Yet, our rate measurements clearly reveal that AOM is not restricted to the uppermost sediments, and that AOM rates peaked at ~17 cm depth, with highest rates of 15 nmol/cm^3^/d (i.e., two orders of magnitude higher than at the sediment surface) (Fig. 1A). The maximum AOM rates observed are comparable to those reported for other freshwater environments (Norði et al. 2013; Segarra et al. 2013, 2015).

Interestingly, the highest AOM activity was observed within sediment layers where our measurements from a parallel core show that sulfate was still available, but at relatively low concentrations (~ 0.1 mmol/L) (Fig. 1B). While this can be taken as indication for AOM coupled to sulfate reduction (Reeburgh 2007), the vertical methane flux and downward diffusion of sulfate suggests an imbalance between the electron donor and its potential electron acceptor in the sediments (−110.5 and 190.9 µmol/m^2^/d for methane and sulfate, respectively). Moreover, a clearly defined sulfate-methane transition zone, as has been commonly described in most diffusive marine settings (Reeburgh 1980, 2007; Iversen and Jørgensen 1985), was not observed at the depth of maximum AOM rates. In fact, relatively high concentrations of both methane and sulfate were found in the surface sediments, where AOM activity was very low, or not detected at all. Furthermore, anaerobic methane turnover continued well below the AOM rate maximum, at depths where sulfate concentrations were almost depleted (~ 0.04 mmol/L). These observations together indicate that AOM was not necessarily limited by the availability of free sulfate within the sediment, and potentially suggest other environmental controls on benthic AOM rates.

The combined geochemical and radiotracer-based rate data imply that AOM in the deeper Lake Cadagno sediments may not depend on free sulfate alone. Anaerobic oxidation of methane coupled to nitrate/nitrite reduction, which has recently been reported for other lake sediments (Deutzmann et al. 2014), seems implausible for Lake Cadagno, as nitrate and nitrite concentrations were below the detection limit (<0.3 µmol/l), both in the euxinic water column and the sediment porewater. While we cannot fully exclude cryptic NOx production by the anaerobic oxidation of ammonium with oxidized metal species (Luther et al. 1997), NO_x_ as an important electron acceptor for AOM in the Lake Cadagno setting seems unlikely. Porewater profiles suggest that the reduction of iron and manganese at, and below, the AOM zone may be involved. Though fermentative/respiratory metal reduction by organotrophs could play a role too in the organic-rich (~15% organic carbon) sediments (Schubert et al. 2011), AOM may be coupled to iron or manganese reduction, as has been suggested elsewhere (Sivan et al. 2011; Crowe et al. 2011; Norði et al. 2013).

### Modes of sedimentary AOM

To further investigate the pathway of AOM in Lake Cadagno sediments, we performed slurry incubation experiments using sediments from three sediment depth segments (Table 1). Segments were chosen based on the methane oxidation rate profiles and the corresponding concentrations of potential electron acceptors (as determined in a separate core). They cover the zone of maximum AOM activity (14-19 cm and 19-24 cm) and the zone below (24-29 cm), where AOM rates were still significant, sulfate concentrations very low, and metal oxide concentrations relatively high (Fig. S1). Moreover, all three segments show the presence of *Candidatus* Methanoperedens, a proven microbial player in AOM (see below). In the first set of experiments (sediments from 14-19 cm and 19-24 cm), we did not detect any methane turnover in killed controls and incubations with 20 mM molybdate, a competitive inhibitor for sulfate reduction (Wilson and Bandurski 1958). In the un-amended live controls (i.e., slurries without additional electron acceptors), at both depths, AOM was slightly elevated relative to killed controls, as indicated by the small amount of excess ^13^CO_2_ that was produced at the end of the incubation period. The low-level AOM might be attributed to ambient substrates remaining in the slurries after their preparation and conditioning (e.g., sulfate). Most strikingly, at both sediment depths, methane oxidation was considerably enhanced upon the addition of either sulfate or MnO_2_ (Fig. 2A and 2B). Excess ^13^CO_2_ production (i.e. methane turnover) was immediately detectable in incubations with added sulfate, whereas a delay of approximately two weeks was observed in our MnO_2_-amended experiments. Though mostly in the incubations with sediments from the 19-24 cm, FeOx-supplemented slurries, similar to the MnO_2_ amendments, showed a stimulation of AOM with a ~2 week delay.

**Figure 2.**
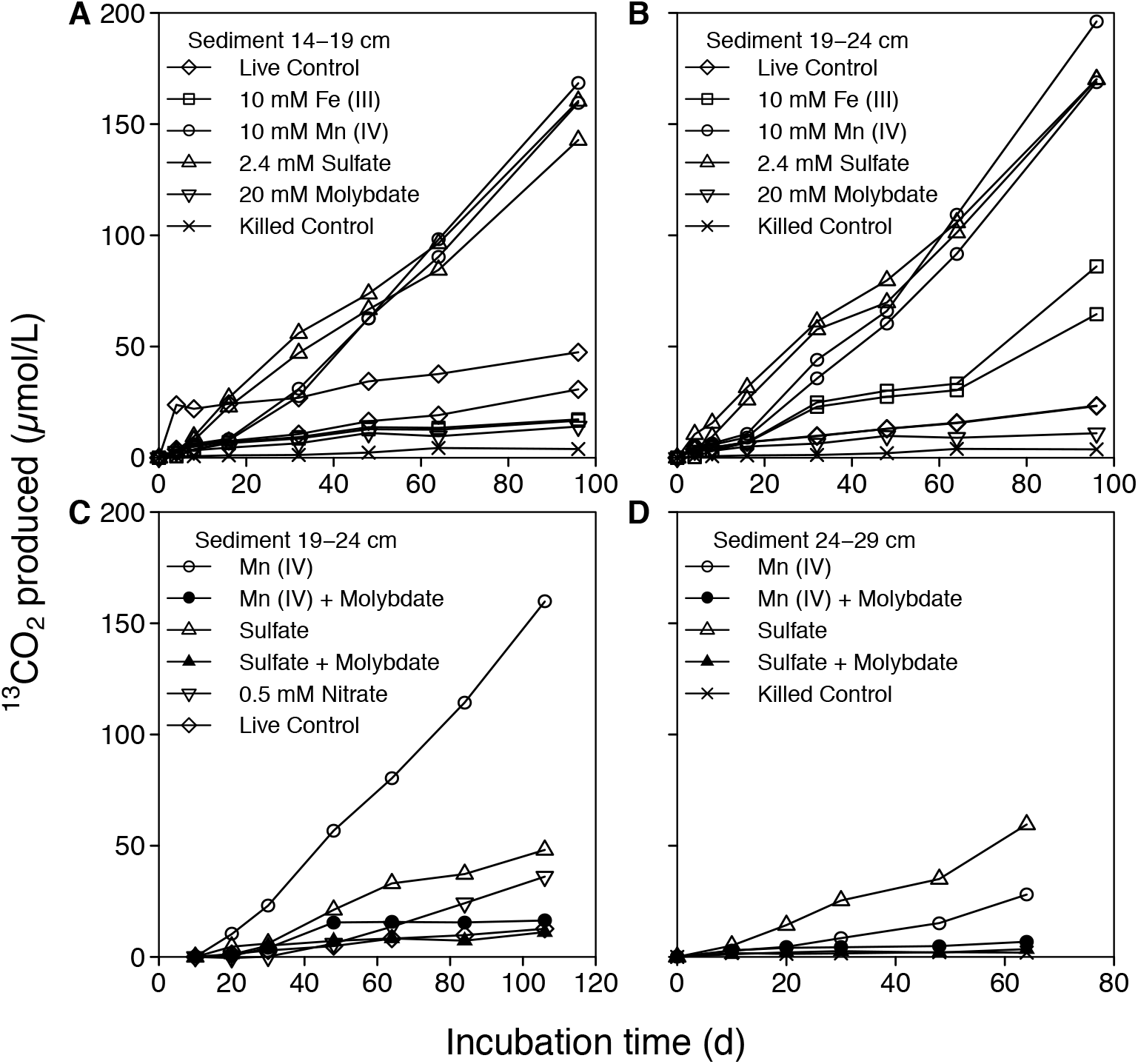
Concentrations of produced ^13^CO_2_ in incubation experiments with ^13^C labeled methane. First set of experiments: slurry incubations with Cadagno sediments from 14-19 cm (A) and 19-24 cm (B) depths, amended with different electron acceptors. Second set of experiments: molybdate inhibition tests in incubations with sulfate and MnO_2_ using sediments from 19-24 cm (C) and 24-29 cm (D), respectively. The effect of nitrate was only tested in slurries from 19-24 cm. In killed-control incubations, autoclaved slurries were used. In live controls no electron acceptors were added.

In the second set of experiments we (re-) amended sediments from 19-24 cm from the first set of experiments, as well as “fresh” sediments from 24-29 cm, with sulfate/MnO_2_/nitrate and/or molybdate (Table 1). This way, we wanted to test whether 1.) AOM was solely and directly driven by sulfate reduction (sulfate-dependent AOM), 2.) whether Mn(IV) reduction was directly coupled to AOM, or 3.) whether the added MnO_2_ fueled a cryptic sulfur or nitrogen cycle, in which alternative electron acceptors (i.e., sulfur intermediates, sulfate or nitrate) were produced through the oxidation of sulfide or ammonium, respectively, with MnO_2_ (Luther et al. 1997; Zopfi et al. 2004; Holmkvist et al. 2011a). During the first 48 days of incubation, no AOM activity was detected in the live control or in incubations with added molybdate. Also thereafter, ^13^CH_4_ oxidation was insignificant in these incubations. In contrast, AOM rates in the nitrate-amended treatment (during the second half of the incubation experiment) were higher compared to the untreated live control, suggesting that, at least under sulfate-depleted conditions, AOM is coupled to denitrification, or that AOM is stimulated by nitrate indirectly. Most obviously, and consistent with the first set of experiments, both sulfate and MnO_2_ boosted ^13^CO_2_ production by AOM compared to the control and the molybdate-addition experiments (Fig. 2C and 2D).

Because AOM was not promoted in incubations with both MnO_2_ and molybdate, in contrast to the MnO_2_-only treatments, we suggest that MnO_2_ was not directly used as electron acceptor during AOM. In the presence of MnO_2_, sulfate can be continuously produced by the oxidation of dissolved sulfide or particulate FeS/FeS_2_ (Yao and Millero 1996; Schippers and Jørgensen 2001), thus fostering sulfate-dependent AOM. Indeed, we observed a net increase in the sulfate concentrations during the incubations with MnO_2_, providing substrate for sulfate-dependent AOM (Fig. S2). In the molybdate-amended experiments, we observed a similar increase in sulfate concentration, which was also likely due to the oxidation of reduced sulfur species with MnO_2_, but AOM was inhibited at the sulfate-reduction step. In the second set of experiments, the enhancing effect of the MnO_2_ addition was approximately three times greater than when sulfate was added to sediments from 19-24 cm (Fig. 2C), while the opposite was observed for sediments from 24-29cm. At this point, we lack an obvious explanation for the observed discrepancy between the MnO_2_ and sulfate-only experiments other than that our observation may leave some scope for true metal-driven AOM in the shallower sediments.

Increasing sulfate concentrations in the incubations with nitrate (Fig. S2) suggests that nitrate might have had a similar effect (i.e., the oxidation of reduced sulfur with nitrate was promoted), but it seemed to stimulate AOM much less than MnO_2_. We conclude that nitrate can, at least in our experiments, like MnO_2_, serve as oxidant for the oxidation of reduced sulfur, producing sulfate that is then available for sulfate-dependent AOM. It is difficult, to explain the weaker effect of nitrate on AOM (i.e., less stimulation compared to the MnO_2_ amendment), if AOM was strictly coupled to sulfate only. More precisely, both the nitrate and MnO_2_ addition resulted in the production of almost equivalent concentrations of sulfate (Fig. S2). This may be taken as additional evidence that MnO_2_ not only affects AOM indirectly through its role in generating sulfate, but also directly by serving as oxidant for true Mn-dependent AOM.

With regards to the effect of FeOx, we expected a stimulation of AOM analogous to that by MnO_2_. However, the overall AOM activity was lower than in the sulfate and MnO_2_ treatments (Fig. 2A and 2B). The weaker ^13^CO_2_ production in the FeOx versus the MnO_2_ treatments is best explained by the fact that sulfate is not a major product of the reaction of FeOx with sulfide (Yao and Millero 1996; Zopfi et al. 2004).

Our incubation results clearly demonstrate that sulfate, added MnOx and FeOx (and potentially nitrate) promoted AOM in the anoxic sediments of Lake Cadagno. In all instances, sulfate appears to be the key electron acceptor used by microorganisms to oxidize methane. We are aware of the limitations with regards to the applicability of high-concentration experimental results to natural environments, and we acknowledge that our combined incubation data leave some scope for true metal-dependent AOM. Yet, we propose that canonical sulfate-dependent AOM is the dominant methane oxidation pathway in the studied sediments. In the upper AOM zone (14-19 cm), where sulfate is replete, AOM is directly coupled to sulfate reduction. In the lower parts of the sediment column (19-29 cm), where free sulfate concentrations are low, sulfate-driven AOM still happens, and is likely maintained by the continuous supply of sulfate produced by the oxidation of reduced sulfur species with metal-oxide phases buried in the sediment (Holmkvist et al. 2011b).

### Lipid biomarker constraints on methane oxidizing microbes

At the end of the slurry incubation period in the first set of experiments with sulfate and manganese oxide, some microbial fatty acids were highly enriched in ^13^C, including monounsaturated C16:1 fatty acids (i.e., C16:1ω7c, C16:1ω7t, C16:1ω5c) and iC17:0 (Fig. 3). High ^13^C-uptake into these lipids most likely by methanotrophic bacteria has been recently observed in other lake sediments (Bar-Or et al. 2017). In contrast, in the live controls, no enrichment of ^13^C was detected in these specific bacterial fatty acids. The extent of ^13^C fatty acid biosynthesis (and thus ^13^CH_4_ uptake) differed both between individual compounds, as well as treatments. The most strongly ^13^C-enriched fatty acid was C16:1ω5c in the 14-19 cm incubation with sulfate (161‰), and in the 19-24 cm incubation with MnO_2_ (307‰), respectively. This fatty acid was previously found in SRB associated to ANME −2 and −3, and to a lesser degree in SRB associated with ANME-1 (Niemann and Elvert 2008). The AOM-associated SRB are known to assimilate the end product of AOM, for example, inorganic carbon (DIC) (Wegener et al. 2008; Kellermann et al. 2012). The observed incorporation of ^13^C can thus be attributed to ^13^CH_4_ oxidation and assimilative uptake of the produced ^13^DIC. Other ^13^C-enriched FA in the treatments with sulfate and/or MnO_2_ (e.g., C16:1ω7c and C16:1ω7t; Fig. 3) were also found in AOM sediments elsewhere, however, they are less useful as a chemotaxonomic marker (Niemann and Elvert 2008). C16:1ω7c is often associated with SRB, but is also present in many other bacteria and eukaryotes. Similarly, iC17:0 has been found in several SRBs (Rütters et al. 2001; Bühring et al. 2006). In any case, the elevated ^13^C/^12^C ratios of the lipids indicate methane-based microbial biosynthesis in our experiments. Strikingly, while we found C16:1ω5c, we did not observe any other fatty acids synthesized by SRBs typically associated with ANMEs, for example the methyl-branched FAs iC15:0 and aiC15:0, or cy-C17:0ω5 and C16:1ω6 (Elvert et al. 2003; Niemann et al. 2006; Niemann and Elvert 2008). Moreover, we did not find any ^13^C-labelled lipids typical for the known archaeal methanotrophs, for example, archaeol and hydroxyarchaeol (Niemann and Elvert 2008), or phytane and biphytane (Bar-Or et al. 2017). We cannot completely rule out that undetectable ^13^C-label incorporation into archaeal lipids during the AOM-incubation experiments may simply be explained by the slow growth of ANME-archaea, with doubling times of several months under laboratory conditions (Nauhaus et al. 2007; Wegener et al. 2008). Yet, we argue that the observed pattern of ^13^C-enriched FAs indicate that AOM in Lake Cadagno is not mediated by any of the known SRB partners. The pattern also does not fit to *Methylomirablis oxyfera* (Kool et al. 2012), the only known bacterium mediating AOM with nitrite (Ettwig et al. 2010).

**Figure 3.**
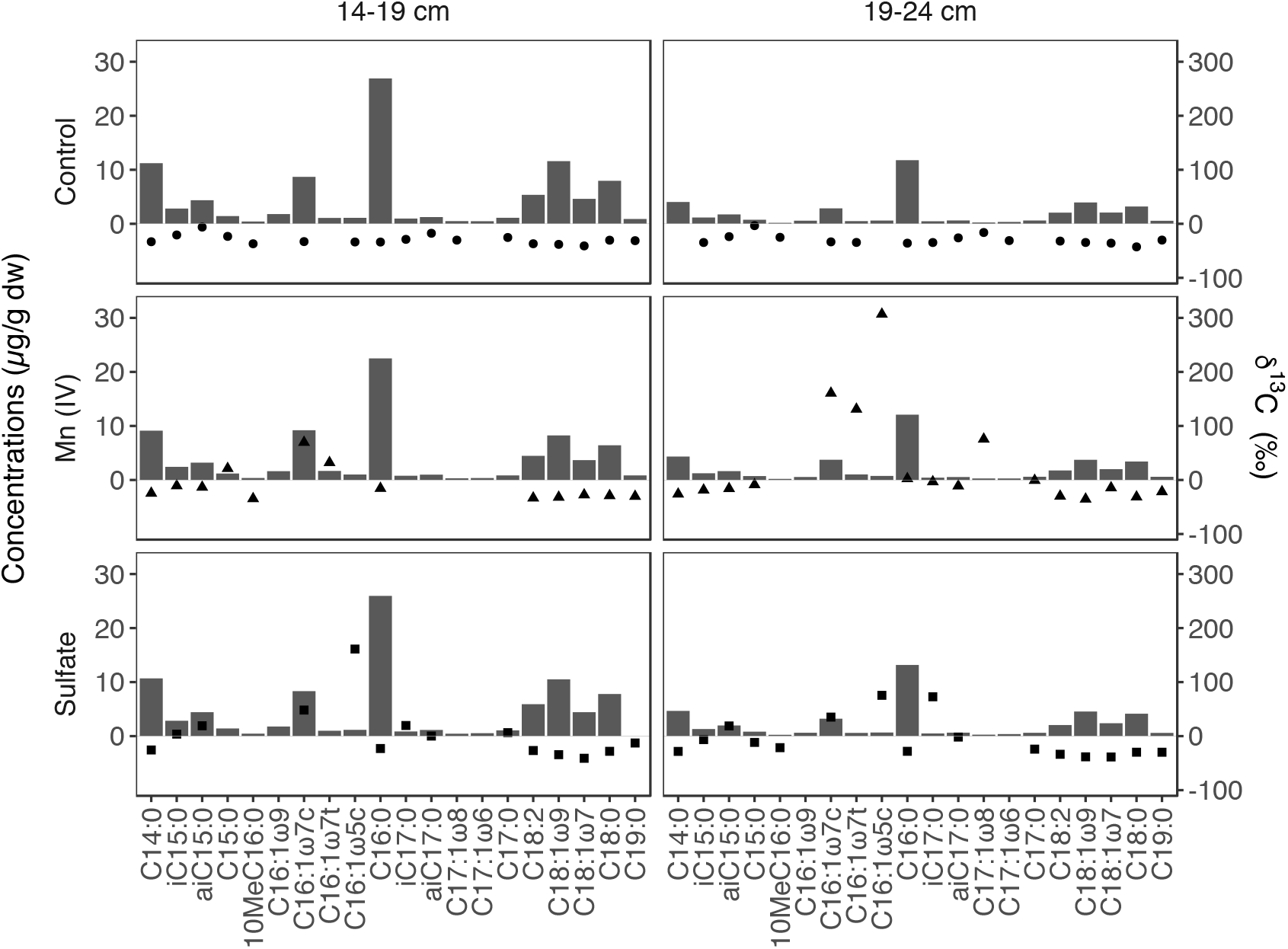
Concentrations (black bars) and compound-specific δ^13^C-values of fatty acids from control experiments (filled circles), samples with MnO_2_ addition (filled triangles) and sulfate (filled squares) of two parallel incubations (sediments from 14-19 cm and 19-24 cm) after 96 days.

### Microorganisms potentially performing AOM

Given the clear evidence of AOM coupled to sulfate reduction in our incubation experiments, we analyzed the sediments for the presence of anaerobic methanotrophic archaea (i.e., ANME-1, −2 and −3) that are typically found in marine sediments in syntrophy with sulfate-reducing bacteria (i.e., Seep-SRB1 and *Desulfobulbus* sp.) (Hinrichs et al. 1999; Boetius et al. 2000; Niemann et al. 2006; Knittel and Boetius 2009). We used primers that match with a large fraction of the deposited sequences of anaerobic methanotrophic archaeal clades (see Supplementary Information, Table S2), but, consistent with our results of the biomarker and gene sequence analyses from the incubation experiments (Fig. 3 and Fig. S3), and with previous 16S rRNA gene analyses in Lake Cadagno sediments (Schubert et al. 2011), we were not able to detect any of the typical ANME-archaea found in marine systems (Knittel and Boetius 2009).

A significant number of 16S rRNA gene sequences that were retrieved at/below the maximum AOM zone (0.3-5.7% of total sequences at >10 cm sediment depth; Fig. 4A), belonged to *Candidatus* Methanoperedens. There is some discrepancy between the vertical distribution of the relative abundance of *Candidatus* Methanoperedens and the AOM rate profile; i.e., the abundance peak was offset by several cm with respect to the maximum AOM rate determined in a separate core. We attribute the apparent offset between the peaks to the heterogeneity of different sediment cores and to lesser degree to artifacts during subsample manipulation. Given that the general shape/quality of the two profiles is essentially equivalent, however, it is reasonable to assume synchronicity, and to link the high methane oxidation rates primarily to this phylotype. The latter was dominated by four amplified sequence variants (ASVs) that showed similar vertical distribution patterns (Fig. S4), and share high sequence similarities within the V4-V5 region of the 16S rRNA gene with recently described *Candidatus* Methanoperedens strains. These are, for example: *Candidatus* Methanoperedens sp. BLZ-1 (Arshad et al. 2015) (98%-100% similarity) and *Candidatus* Methanoperedens nitroreducens ANME-2d (Haroon et al. 2013) (97%-99% similarity) in bioreactors, and *Candidatus* Methanoperedens nitroreducens Vercelli in paddy field soils (Vaksmaa et al. 2017), which are capable of coupling methane oxidation to nitrate reduction (Haroon et al. 2013). However, recent RNA stable isotope probing results suggest that archaea of this clade may also perform AOM coupled to iron and/or sulfate reduction in the iron-rich but sulfate-poor sediments of Lake Ørn (Denmark) (Weber et al. 2017). The ANME-2d sequences in these sediments share >98% sequence similarity with the Methanoperedenaceae phylotypes found in Lake Cadagno (Fig. S5). Moreover, there is genomic evidence for the presence of numerous multiheme c-type cytochromes in Methanoperedens-like archaea, suggesting that they can transfer electrons to a broad range of electron acceptors (Arshad et al. 2015; McGlynn et al. 2015), as shown experimentally for Fe(III)- and Mn(IV)-dependent AOM (Ettwig et al. 2016). Multiheme c-type cytochromes in ANME-1 archaea have also been shown to facilitate the electron transfer to syntrophic partner organisms such as SRB (Wegener et al. 2015).

**Figure 4.**
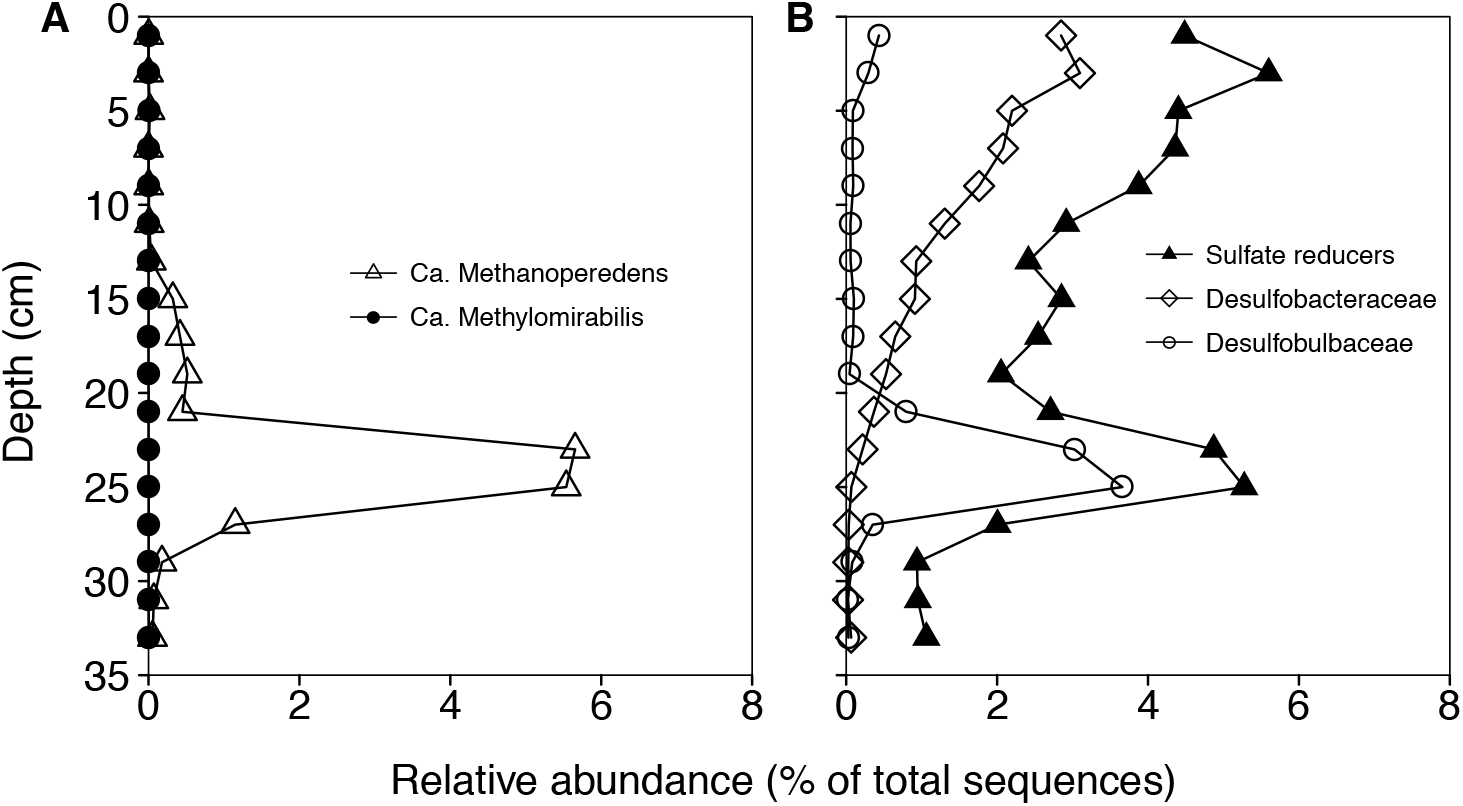
Depth profiles of relative abundances of (A) *Candidatus* Methanoperedens and *Candidatus* Methylomirabilis and (B) sulfate-reducing bacteria (SRBs) in the sediments of Lake Cadagno. Among the SRBs, representatives of Desulfobulbaceae and Desulfobacteraceae have been shown to form consortia with anaerobic methane oxidizing archaea. Data are based on read abundances of 16S rRNA gene sequences.

In Lake Cadagno sediments, we also found sequences of sulfate reducing bacteria, including the SEEP-SRB1 cluster and members of the Desulfobulbus group, which have been shown to be associated to ANME-1, −2 and −3, respectively, as bacterial partners in marine settings (Knittel and Boetius 2009). The relative abundances of SRBs decreased with sediment depth, in parallel with decreasing sulfate concentrations, but then increased again at about 24 cm depth (Fig. 4B). This secondary maximum was due to a local enrichment of Desulfobulbaceae, which were dominated by a single uncultured representative of this family (Figs. S4, S5, S6). The correspondence of high abundances of Desulfobulbaceae and the *Candidatus* Methanoperedens peak at the same sediment depth (Fig. 4B and Fig. S7), together with the lipid-SIP results, suggests that the anaerobic methane-oxidizing archaea and sulfate-reducing bacteria detected in Lake Cadagno sediments are interdependent. The exact nature of the syntrophic interaction (e.g., formation of consortia or an indirect association of SRB and AOM) awaits further investigation. We note, however, that this interdependence is most likely facultative, as one of the amplified sequence variants of *Ca.* Methanoperedens (ZOTU202) showed a second abundance peak at a depth (13-19 cm), where no Desulfobulbaceae partner sequences were detected (Fig. S4A; Fig. 4B). This is consistent with our incubation experiments, and suggests that at least this strain of *Ca.* Methanoperedens can perform AOM independently, provided a suitable electron acceptor is present.

## Concluding remarks

In the present study, patterns of AOM activity, pathways and microbial diversity were investigated in the sediments of euxinic Lake Cadagno. We present clear evidence that microorganisms performed anaerobic oxidation of methane coupled to sulfate reduction below 10 cm sediment depth, with relatively high AOM rates even at depths where sulfate concentrations are relatively low. Incubation experiments show that the addition of sulfate, manganese, iron, and/or nitrate promotes AOM. While there is some evidence for metal oxide-dependent AOM, we argue that the stimulation of AOM by the non-sulfate oxidants was mostly indirect. Sulfate-dependent methane oxidation was fueled by continuous (and at greater depths, cryptic) sulfate production by the oxidation of reduced sulfur compounds with metal oxides. Our microbial community analysis revealed that AOM was driven by uncultured archaea of the candidate genus Methanoperedens. The parallel depth distribution of the abundances of *Candidatus* Methanoperedens and potential sulfate-reducing ANME partners in the sediment zone where high AOM rates were observed suggests that methane oxidation is performed in archaeal-bacterial association. However, it cannot be excluded that the methane oxidizing archaea are able to perform sulfate-dependent AOM by themselves. The coupling of AOM to sulfate reduction by novel Methanoperedenaceae (and the possible disguise as Mn-/Fe-dependent methanotrophs) not only expands our understanding of this biogeochemically significant group and their potential for metabolic versatility, but also has broad implications for future AOM investigations in freshwater environments, where sulfate concentrations are low and metal (Mn, Fe) concentrations are often high. Here, *Candidatus* Methanoperedens may represent important sentinels of methane emission to the atmosphere, taking over a similar ecological role as ANME-1, −2 and −3 in marine sediments.

## Supporting information

Manuscript

## Acknowledgments

We thank Thomas Kuhn for his excellent support in the laboratory and Jean-Claude Walser at the Genetic Diversity Centre (ETH, Zürich) for bioinformatic support. We are grateful to Judith Kobler Waldis for measuring iron and manganese concentrations and thank Ruth Strunk and Nikolaus Kuhn for their help with DIC concentrations measurements. Carsten Schubert and Serge Robert (EAWAG) are especially acknowledged for providing access to laboratory infrastructure and assistance with methane isotope measurements. We also thank Jana Tischer and Maciej Bartosiewicz for their assistance during field campaigns, and the personnel of the Piora Centro Biologia Alpina for providing access to the sampling platform. This research was supported by the China Scholarship Council (CSC). Additional funding came from the Department of Environmental Sciences of the University Basel.

## Author contributions

MFL, JZ and HN conceived the research. GS performed all the experiments with support from JZ. GS, MFL, JZ and HN performed data analyses and interpretation. HY and LS assisted in the interpretation of lipid biomarker data. GS prepared the manuscript with support from MFL, JZ, and HN.

## References

Andrews, S. 2018. FastQC: a quality control tool for high throughput sequence data.

Arshad, A., D. R. Speth, R. M. De Graaf, H. J. M. Op den Camp, M. S. M. Jetten, and C. U. Welte. 2015. A metagenomics-based metabolic model of nitrate-dependent anaerobic oxidation of methane by Methanoperedens-like archaea. Front. Microbiol. 6: 1–14. doi:10.3389/fmicb.2015.01423

Bar-Or, I., M. Elvert, W. Eckert, A. Kushmaro, H. Vigderovich, Q. Zhu, E. Ben-Dov, and O. Sivan. 2017. Iron-coupled anaerobic oxidation of methane performed by a mixed bacterial-archaeal community based on poorly reactive minerals. Environ. Sci. Technol. 51: 12293–12301. doi:10.1021/acs.est.7b03126

Bastviken, D., J. Ejlertsson, and L. Tranvik. 2002. Measurement of methane oxidation in lakes: A comparison of methods. Environ. Sci. Technol. 36: 3354–3361. doi:10.1021/es010311p

Beal, E. J., C. H. House, and V. J. Orphan. 2009. Manganese- and Iron-Dependent Marine Methane Oxidation. Science. 325: 184–187. doi:10.1126/science.1169984

Blees, J., H. Niemann, C. B. Wenk, and others. 2014. Micro-aerobic bacterial methane oxidation in the chemocline and anoxic water column of deep south-Alpine Lake Lugano (Switzerland). Limnol Ocean. 59: 311–324. doi:10.4319/lo.2014.59.2.0311

Boetius, A., K. Ravenschlag, C. J. Schubert, and others. 2000. A marine microbial consortium apparently mediating anaerobic oxidation of methane. Nature 407: 623–626. doi:10.1038/35036572

Boudreau, B. P. 1997. Diagenetic models and their implementation: Modelling transport and reactions in aquatic sediments, Springer-Verlag.

Bühring, S. I., N. Lampadariou, L. Moodley, A. Tselepides, and U. Witte. 2006. Benthic microbial and whole-community responses to different amounts of ^13^C-enriched algae: In situ experiments in the deep Cretan Sea (Eastern Mediterranean). Limnol Ocean. 51: 157–165.

Cai, C., A. O. Leu, G.-J. Xie, and others. 2018. A methanotrophic archaeon couples anaerobic oxidation of methane to Fe(III) reduction. ISME J. 12: 1929–1939. doi:10.1038/s41396-018-0109-x

Cline, J. D. 1969. Spectrophotometric determination of hydrogen sulfide in natural waters. Limnol Ocean. 14: 454–458. doi:10.4319/lo.1969.14.3.0454

Crowe, S. A., S. Katsev, K. Leslie, and others. 2011. The methane cycle in ferruginous Lake Matano. Geobiology 9: 61–78. doi:10.1111/j.1472-4669.2010.00257.x

Deutzmann, J. S., P. Stief, J. Brandes, and B. Schink. 2014. Anaerobic methane oxidation coupled to denitrification is the dominant methane sink in a deep lake. Proc. Natl. Acad. Sci. 111: 18273–18278. doi:10.1073/pnas.1411617111

Edgar, R. 2016. SINTAX: a simple non-Bayesian taxonomy classifier for 16S and ITS sequences. bioRxiv. doi:https://doi.org/10.1101/074161

Edgar, R. C. 2013. UPARSE: highly accurate OTU sequences from microbial amplicon reads. Nat. Methods 10: 996–1000. doi:10.1038/nmeth.2604

Egger, M., O. Rasigraf, C. J. Sapart, and others. 2015. Iron-mediated anaerobic oxidation of methane in brackish coastal sediments. Environ. Sci. Technol. 49: 277–283. doi:10.1021/es503663z

Elvert, M., A. Boetius, K. Knittel, and B. B. Jørgensen. 2003. Characterization of specific membrane fatty acids as chemotaxonomic markers for sulfate-reducing bacteria involved in anaerobic oxidation of methane. Geomicrobiol. J. 20: 403–419. doi:10.1080/01490450303894

Ettwig, K. F., T. Van Alen, K. T. Van De Pas-Schoonen, M. S. M. Jetten, and M. Strous. 2009. Enrichment and molecular detection of denitrifying methanotrophic bacteria of the NC10 phylum. Appl Env. Microbiol 75: 3656–3662. doi:10.1128/AEM.00067-09

Ettwig, K. F., M. K. Butler, D. Le Paslier, and others. 2010. Nitrite-driven anaerobic methane oxidation by oxygenic bacteria. Nature 464: 543–548. doi:10.1038/nature08883

Ettwig, K. F., B. Zhu, D. Speth, J. T. Keltjens, M. S. M. Jetten, and B. Kartal. 2016. Archaea catalyze iron-dependent anaerobic oxidation of methane. Proc. Natl. Acad. Sci. 113: 12792–12796. doi:10.1073/pnas.1609534113

Graf, J. S., M. J. Mayr, H. K. Marchant, and others. 2018. Bloom of a denitrifying methanotroph, “ Candidatus Methylomirabilis limnetica”, in a deep stratified lake. Environ. Microbiol. 20: 2598–2614. doi:10.1111/1462-2920.14285

Hansel, C. M., C. J. Lentini, Y. Tang, D. T. Johnston, S. D. Wankel, and P. M. Jardine. 2015. Dominance of sulfur-fueled iron oxide reduction in low-sulfate freshwater sediments. ISME J. 9: 2400–2412. doi:10.1038/ismej.2015.50

Haroon, M. F., S. Hu, Y. Shi, M. Imelfort, J. Keller, P. Hugenholtz, Z. Yuan, and G. W. Tyson. 2013. Anaerobic oxidation of methane coupled to nitrate reduction in a novel archaeal lineage. Nature 500: 567–70. doi:10.1038/nature12375

Hinrichs, K. U., J. M. Hayes, S. P. Sylva, P. G. Brewer, and E. F. DeLong. 1999. Methane-consuming archaebacteria in marine sediments. Nature 398: 802–805. doi:10.1038/19751

Holmkvist, L., T. G. Ferdelman, and B. Barker. 2011a. A cryptic sulfur cycle driven by iron in the methane zone of marine sediment (Aarhus Bay, Denmark). Geochim. Cosmochim. Acta 75: 3581–3599. doi:10.1016/j.gca.2011.03.033

Holmkvist, L., A. Kamyshny, C. Vogt, K. Vamvakopoulos, T. G. Ferdelman, and B. B. Jørgensen. 2011b. Sulfate reduction below the sulfate-methane transition in Black Sea sediments. Deep. Res. Part I Oceanogr. Res. Pap. 58: 493–504. doi:10.1016/j.dsr.2011.02.009

Iversen, N., and B. B. Jørgensen. 1985. Anaerobic methane oxidation rates at the sulfate-methane transition in marine sediments from Kattegat and Skagerrak (Denmark). Limnol Ocean. 30: 944–955.

Kellermann, M. Y., G. Wegener, M. Elvert, and others. 2012. Autotrophy as a predominant mode of carbon fixation in anaerobic methane-oxidizing microbial communities. Proc. Natl. Acad. Sci. 109: 19321–19326. doi:10.1073/pnas.1208795109

Knittel, K., and A. Boetius. 2009. Anaerobic oxidation of methane: progress with an unknown process. Annu. Rev. Microbiol. 63: 311–334. doi:10.1146/annurev.micro.61.080706.093130

Kool, D. M., B. Zhu, W. I. C. Rijpstra, M. S. M. Jetten, K. F. Ettwig, and J. S. Sinninghe Damsté. 2012. Rare branched fatty acids characterize the lipid composition of the intra-aerobic methane oxidizer “Candidatus Methylomirabilis oxyfera.” Appl Env. Microbiol 78: 8650–8656. doi:10.1128/AEM.02099-12

Luther, G. W., B. Sundby, B. L. Lewis, P. J. Brendel, and N. Silverberg. 1997. Interactions of manganese with the nitrogen cycle: Alternative pathways to dinitrogen. Geochim. Cosmochim. Acta 61: 4043–4052. doi:10.1016/S0016-7037(97)00239-1

Magoč, T., and S. L. Salzberg. 2011. FLASH: fast length adjustment of short reads to improve genome assemblies. Bioinformatics 27: 2957–2963. doi:10.1093/bioinformatics/btr507

Martinez-Cruz, K., A. Sepulveda-Jauregui, P. Casper, K. W. Anthony, K. A. Smemo, and F. Thalasso. 2018. Ubiquitous and significant anaerobic oxidation of methane in freshwater lake sediments. Water Res. 144: 332–340. doi:10.1016/j.watres.2018.07.053

McGlynn, S. E., G. L. Chadwick, C. P. Kempes, and V. J. Orphan. 2015. Single cell activity reveals direct electron transfer in methanotrophic consortia. Nature 526: 531–535. doi:10.1038/nature15512

McMurdie, P. J., and S. Holmes. 2013. Phyloseq: An R Package for Reproducible Interactive Analysis and Graphics of Microbiome Census Data. PLoS One 8: 1–11. doi:10.1371/journal.pone.0061217

Milucka, J., T. G. Ferdelman, L. Polerecky, and others. 2012. Zero-valent sulphur is a key intermediate in marine methane oxidation. Nature 491: 541–6. doi:10.1038/nature11656

Moss, C. W., and M. A. Lambert-fair. 1989. Location of double bonds in monounsaturated fatty acids of Campylobacter cryaerophila with dimethyl disulfide derivatives and combined gas chromatography-mass spectrometry. J. Clin. Microbiol. 27: 1467–1470.

Nauhaus, K., M. Albrecht, M. Elvert, A. Boetius, and F. Widdel. 2007. In vitro cell growth of marine archaeal-bacterial consortia during anaerobic oxidation of methane with sulfate. Environ. Microbiol. 9: 187–196. doi:10.1111/j.1462-2920.2006.01127.x

Nichols, P. D., J. B. Guckert, and D. C. White. 1986. Determination of monosaturated fatty acid double-bond position and geometry for microbial monocultures and complex consortia by capillary GC-MS of their dimethyl disulphide adducts. J. Microbiol. Methods 5: 49–55. doi:https://doi.org/10.1016/0167-7012(86)90023-0

Niemann, H., and M. Elvert. 2008. Diagnostic lipid biomarker and stable carbon isotope signatures of microbial communities mediating the anaerobic oxidation of methane with sulphate. Org. Geochem. 39: 1668–1677. doi:10.1016/j.orggeochem.2007.11.003

Niemann, H., M. Elvert, M. Hovland, and others. 2005. Methane emission and consumption at a North Sea gas seep (Tommeliten area). Biogeosciences 2: 335–351. doi:10.5194/bg-2-335-2005

Niemann, H., T. Lösekann, D. de Beer, and others. 2006. Novel microbial communities of the Haakon Mosby mud volcano and their role as a methane sink. Nature 443: 854–858. doi:10.1038/nature05227

Niemann, H., L. Steinle, J. Blees, I. Bussmann, T. Treude, S. Krause, M. Elvert, and M. F. Lehmann. 2015. Toxic effects of lab-grade butyl rubber stoppers on aerobic methane oxidation. Limnol Ocean. Methods 13: 40–52. doi:10.1002/lom3.10005

Norði, K. Á., B. Thamdrup, and C. J. Schubert. 2013. Anaerobic oxidation of methane in an iron-rich Danish freshwater lake sediment. Limnol Ocean. 58: 546–554. doi:10.4319/lo.2013.58.2.0546

Orphan, V. J., C. H. House, K.-U. Hinrichs, K. D. McKeegan, and E. F. DeLong. 2002. Multiple archaeal groups mediate methane oxidation in anoxic cold seep sediments. Proc. Natl. Acad. Sci. 99: 7663–7668. doi:10.1073/pnas.072210299

Oswald, K., J. Milucka, A. Brand, S. Littmann, B. Wehrli, M. M. M. Kuypers, and C. J. Schubert. 2015. Light-dependent aerobic methane oxidation reduces methane emissions from seasonally stratified lakes. PLoS One 10: 1–22. doi:10.1371/journal.pone.0132574

Parada, A. E., D. M. Needham, and J. A. Fuhrman. 2016. Every base matters: Assessing small subunit rRNA primers for marine microbiomes with mock communities, time series and global field samples. Environ. Microbiol. 18: 1403–1414. doi:10.1111/1462-2920.13023

Quast, C., E. Pruesse, P. Yilmaz, J. Gerken, T. Schweer, P. Yarza, J. Peplies, and F. O. Glöckner. 2013. The SILVA ribosomal RNA gene database project: improved data processing and web-based tools. Nucleic Acids Res. 41: 590–596. doi:10.1093/nar/gks1219

Raghoebarsing, A. a, A. Pol, K. T. van de Pas-Schoonen, and others. 2006. A microbial consortium couples anaerobic methane oxidation to denitrification. Nature 440: 918–921. doi:10.1038/nature04617

Reeburgh, W. 2007. Oceanic methane biogeochemistry. Am. Chem. Soc. 107: 486–513. doi:10.1021/cr050362v

Reeburgh, W. S. 1980. Anaerobic methane oxidation: Rate depth distributions in Skan Bay sediments. Earth Planet. Sci. Lett. 47: 345–352. doi:10.1016/j.jom.2015.11.003

Rütters, H., H. Sass, H. Cypionka, and J. Rullkötter. 2001. Monoalkylether phospholipids in the sulfate-reducing bacteria Desulfosarcina variabilis and Desulforhabdus amnigenus. Arch. Microbiol. 176: 435–442. doi:10.1007/s002030100343

Schippers, A., and B. B. Jørgensen. 2001. Oxidation of pyrite and iron sul de by manganese dioxide in marine sediments. Geochim. Cosmochim. Acta 65: 915–922.

Schmieder, R., and R. Edwards. 2011. Quality control and preprocessing of metagenomic datasets. Bioinformatics 27: 863–864. doi:10.1093/bioinformatics/btr026

Schubert, C. J., F. Vazquez, T. Lösekann-Behrens, K. Knittel, M. Tonolla, and A. Boetius. 2011. Evidence for anaerobic oxidation of methane in sediments of a freshwater system (Lago di Cadagno). FEMS Microbiol. Ecol. 76: 26–38. doi:10.1111/j.1574-6941.2010.01036.x

Segarra, K. E. A., C. Comerford, J. Slaughter, and S. B. Joye. 2013. Impact of electron acceptor availability on the anaerobic oxidation of methane in coastal freshwater and brackish wetland sediments. Geochim. Cosmochim. Acta 115: 15–30. doi:10.1016/j.gca.2013.03.029

Segarra, K. E. a., F. Schubotz, V. Samarkin, M. Y. Yoshinaga, K.-U. Hinrichs, and S. B. Joye. 2015. High rates of anaerobic methane oxidation in freshwater wetlands reduce potential atmospheric methane emissions. Nat Commun 6: 1–8. doi:10.1038/ncomms8477

Sivan, O., M. Adler, A. Pearson, F. Gelman, I. Bar-Or, S. G. John, and W. Eckert. 2011. Geochemical evidence for iron-mediated anaerobic oxidation of methane. Limnol Ocean. 56: 1536–1544. doi:10.4319/lo.2011.56.4.1536

Steinle, L., M. Schmidt, L. Bryant, and others. 2016. Linked sediment and water-column methanotrophy at a man-made gas blowout in the North Sea: Implications for methane budgeting in seasonally stratified shallow seas. Limnol Ocean. 61: S367–S386. doi:10.1002/lno.10388

Stookey, L. L. 1970. Ferrozine-A new spectrophotometric reagent for iron. Anal. Chem. 42: 779–781. doi:10.1021/ac60289a016

Timmers, P. H. A., H. C. A. Widjaja-Greefkes, J. Ramiro-Garcia, C. M. Plugge, and A. J. M. Stams. 2015. Growth and activity of ANME clades with different sulfate and sulfide concentrations in the presence of methane. Front. Microbiol. 6. doi:10.3389/fmicb.2015.00988

Timmers, P. H., D. A. Suarez-Zuluaga, M. van Rossem, M. Diender, A. J. Stams, and C. M. Plugge. 2016. Anaerobic oxidation of methane associated with sulfate reduction in a natural freshwater gas source. ISME J. 10: 1400–1412. doi:10.1038/ismej.2015.213

Vaksmaa, A., S. Guerrero-Cruz, T. A. van Alen, G. Cremers, K. F. Ettwig, C. Lüke, and M. S. M. Jetten. 2017. Enrichment of anaerobic nitrate-dependent methanotrophic ‘*Candidatus* Methanoperedens nitroreducens’ archaea from an Italian paddy field soil. Appl. Microbiol. Biotechnol. 101: 7075–7084. doi:10.1007/s00253-017-8416-0

Versantvoort, W., S. Guerrero-Cruz, D. R. Speth, and others. 2018. Comparative genomics of *Candidatus* Methylomirabilis species and description of Ca. Methylomirabilis lanthanidiphila. Front. Microbiol. 9: 1–10. doi:10.3389/fmicb.2018.01672

Wang, F.-P., Y. Zhang, Y. Chen, and others. 2014. Methanotrophic archaea possessing diverging methane-oxidizing and electron-transporting pathways. ISME J. 8: 1069–1078. doi:10.1038/ismej.2013.212

Weber, H. S., K. S. Habicht, and B. Thamdrup. 2017. Anaerobic methanotrophic archaea of the ANME-2d cluster are active in a low-sulfate, iron-rich freshwater sediment. Front. Microbiol. 8: 1–13. doi:10.3389/fmicb.2017.00619

Wegener, G., V. Krukenberg, D. Riedel, H. E. Tegetmeyer, and A. Boetius. 2015. Intercellular wiring enables electron transfer between methanotrophic archaea and bacteria. Nature 526: 587–590. doi:10.1038/nature15733

Wegener, G., H. Niemann, M. Elvert, K. U. Hinrichs, and A. Boetius. 2008. Assimilation of methane and inorganic carbon by microbial communities mediating the anaerobic oxidation of methane. Environ. Microbiol. 10: 2287–2298. doi:10.1111/j.1462-2920.2008.01653.x

Wilson, L. G., and R. S. Bandurski. 1958. Enzymatic reactions involving sulfate, sulfite, selenate and molybdate. J. Biol. Chem. 233: 975–981.

Yao, W., and F. J. Millero. 1996. Oxidation of hydrogen sulfide by hydrous Fe (III) oxides in seawater. Mar. Chem. 52: 1–16. doi:10.1016/0304-4203(95)00072-0

Zopfi, J., T. G. Ferdelman, and H. Fossing. 2004. Distribution and fate of sulfur intermediates - sulfite, tetrathionate, thiosulfate and elemental sulfur - in marine sediments. Spec. Pap. Geol. Soc. Am. 379: 17–34. doi:10.1130/0-8137-2379-5

