## Supplementary material for "Manganese/iron-supported sulfate-dependent anaerobic oxidation of methane by archaea in lake sediments": Manuscript

### Supplementary information on methods

Preparation of Fe(III)- and Mn(IV)-oxides.
Two-line Ferrihydrite ($Fe_5O_8H \cdot H_2O$, hereafter "FeOx") for $^{13}CH_4$ incubation experiments was produced according to Schwertmann and Cornell (Schwertmann and Cornell 2000).
Reaction: $5FeCl_3 + 15NaOH \rightarrow Fe_5O_8H \cdot H_2O + 15NaCl + 6H_2O$

1. In a large beaker, 27g $FeCl_3$ was dissolved in about 300 mL distilled water, and NaOH solution (20 % w/w) was added until pH 7 is reached.
2. The suspension was stirred for around 30 min and the FeOx was washed (by decanting/resuspending) several times with distilled water until excess NaCl was removed. FeOx was kept as an aqueous slurry under $N_2$ atmosphere in glass bottles (Schott) with thick butyl rubber septa.

Preparation of $MnO_2$ (Vernadite) according to Kostka and Nealson (1998).

Reaction: $2KMnO_4 + 3MnCl_2 + 2H_2O \rightarrow 5MnO_2 + 4H^+ + 2K^+ + 6Cl^-$

1. 5.92g $KMnO_4$ was dissolved in 148.2 mL distilled water, heated to 90 °C, then 7.5ml 5N NaOH was added.
2. 13 g $MnCl_2 \cdot 4H_2O$ was dissolved in 60 ml distilled water and then slowly added to the alkaline $KMnO_4$ solution.
3. Heating was stopped, and after 30 minutes of continued stirring at room temperature, the suspension was washed several times to remove excess salts and acidity. $MnO_2$ was kept as an aqueous slurry under $N_2$ atmosphere in glass bottles (Schott) with thick butyl rubber septa.

### Supplementary Figures

Geochemical data

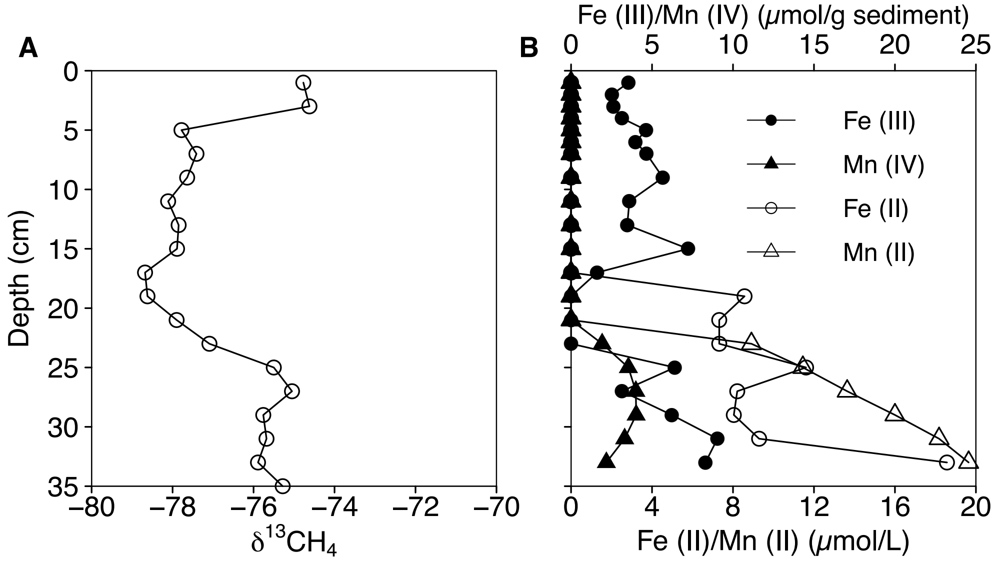


**Figure S1.** Depth profiles of (A) $\delta^{13}CH_4$ (in ‰ vs. V-PDB) in the sediments of Lake Cadagno and (B) concentrations of dissolved Fe(II), Mn(II), HCl-extractable Fe(III)) and solid Mn-phases. Fe(II)/Mn(II) concentrations in the porewater likely included metal complexes or colloids that passed through the 0.45 μm filter pores, as they should not coexist freely with sulfide at the depth interval of 19-25 cm.

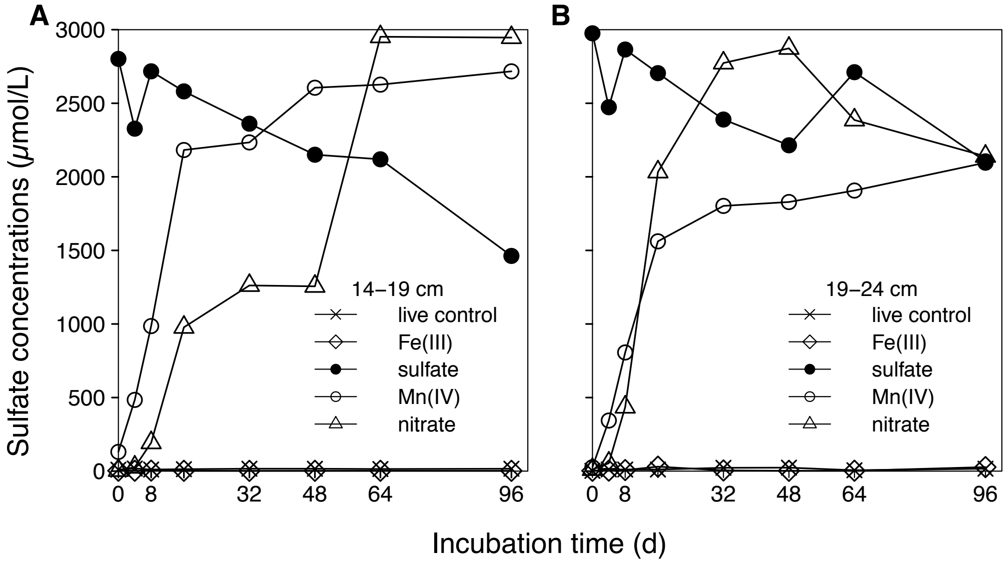


**Figure S2.** Sulfate concentrations with incubation time in slurry incubations (first set of experiments) using sediments from (A) 14-19 cm and (B) 19-24 cm depth in Lake Cadagno. Different symbols represent the different electron acceptors added.

### Molecular data

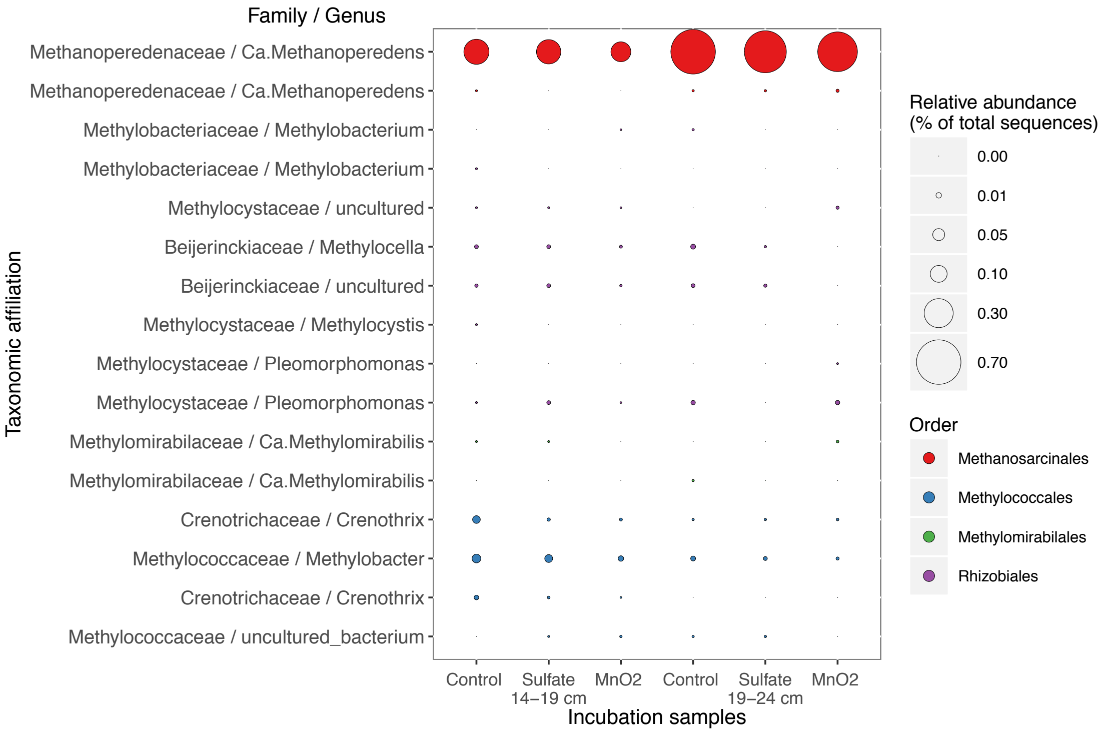


**Figure S3.** Relative abundances of aerobic and anaerobic methanotrophs detected in incubation samples with sulfate and manganese oxide, and in control experiments after 96 days. Consistent with the down-core molecular data, incubations were dominated by 16S rRNA gene sequences affiliated to *Candidatus* Methanoperedens, implying that AOM in the Lake Cadagno sediments is primarily driven by this phylotype. The relative abundance of *Ca.* Methanoperedens appeared to decrease during the incubation, which was most likely due to growth of other organisms (e.g., SRBs upon sulfate addition).

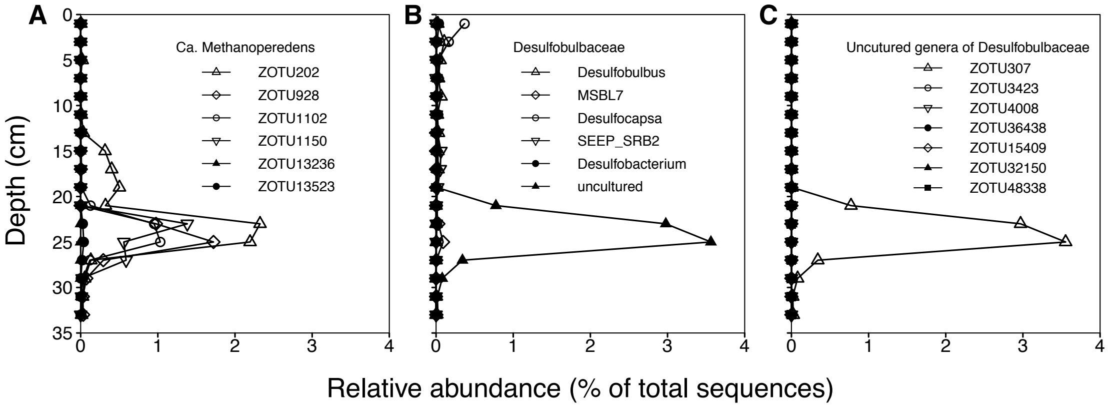


**Figure S4.** (A) Relative abundances of the amplicon sequence variants (i.e. zero-radius OTU, ZOTU) forming the *Ca*. Methanoperedens phylotype in the sediments of Lake Cadagno. The four dominant ZOTUs peaked at the same depth as uncultured Desulfobulbaceae (B), a family previously shown to be associated with anaerobic methanotrophs in marine environments. (C) The depth distribution of all detected Desulfobulbaceae ZOTUs reveals that this family was dominated by a single amplified sequence variant (ZOTU307). Interestingly, *Ca.* Methanoperedens ZOTU202 showed a second abundance peak at a depth (13-19 cm) where ZOTU307 was not detected, suggesting that this strain may also be able to perform AOM independently in presence of a suitable electron acceptor. Data are based on read abundances of 16S rRNA gene sequences.

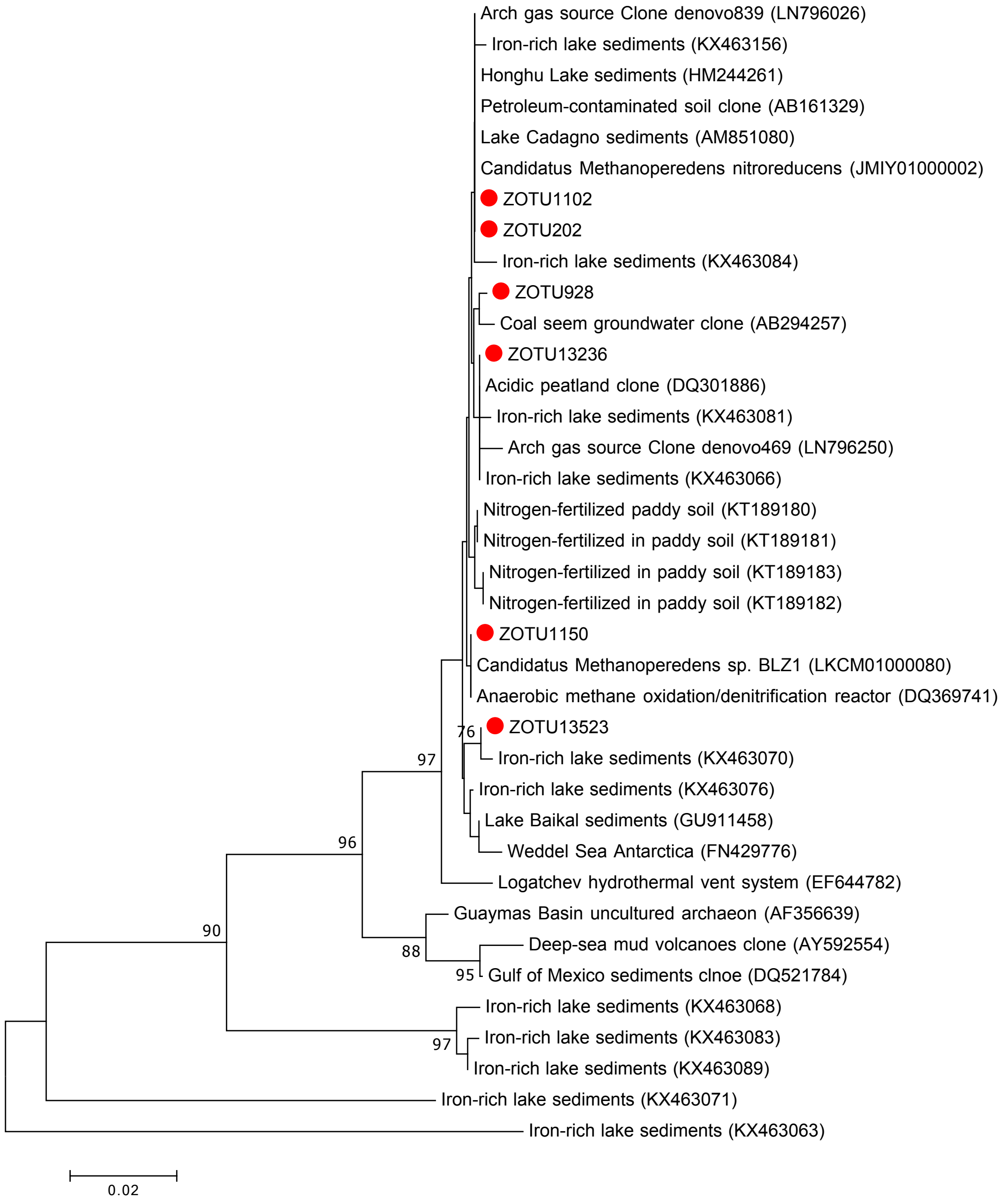


**Figure S5.** Neighbor-joining tree showing representative 16S rRNA gene sequences of Methanoperedenaceae including the amplified sequence variants (ZOTU) retrieved form Lake Cadagno sediments (red circles) and closely related sequences from different environments. The tree was constructed using Maximum Composite Likelihood correction and complete gap deletion (Kumar et al. 2016). Bootstrap values >65% for 1000 re-samplings are shown and the scale bar represents 2% estimated sequence divergence.

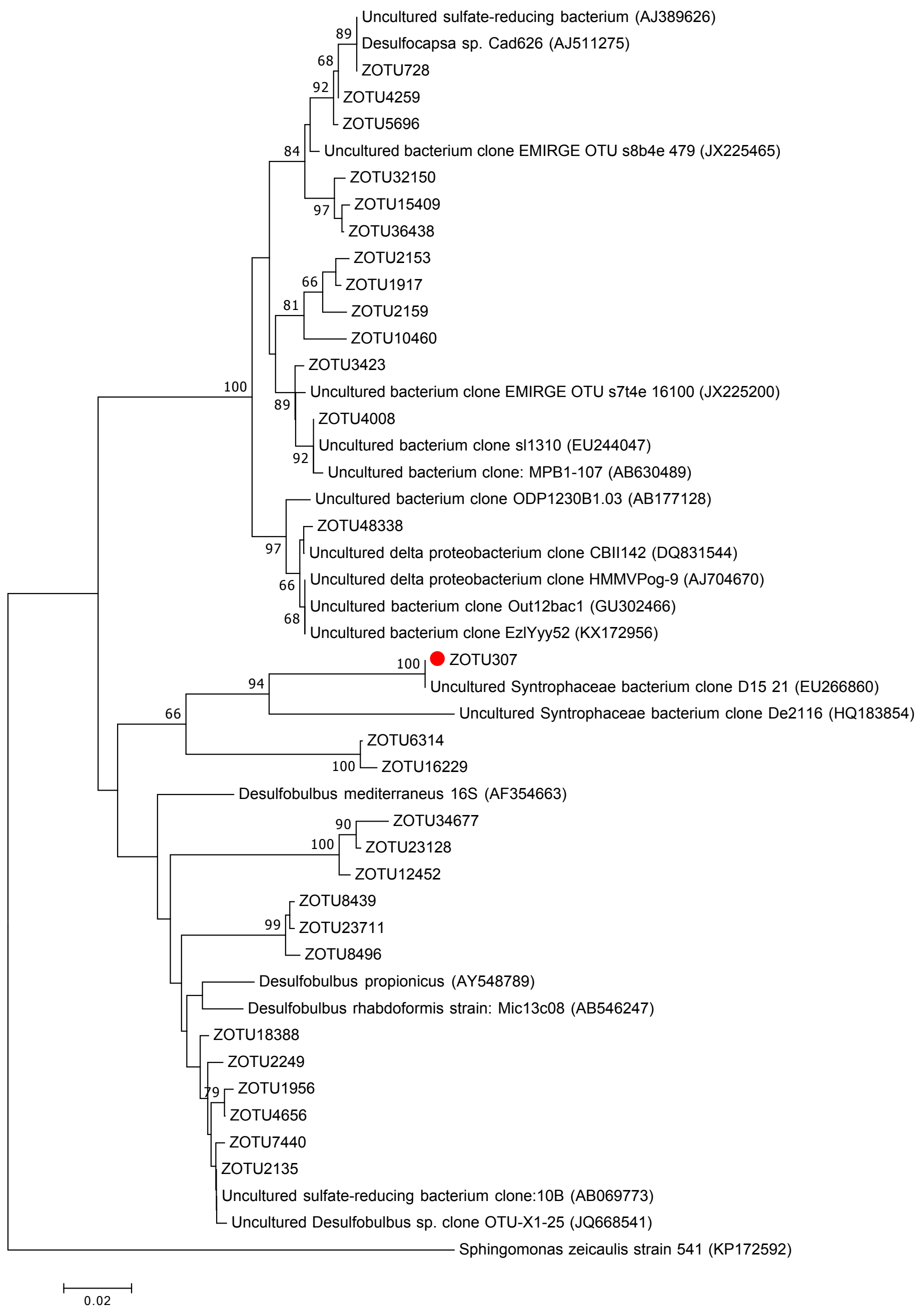


**Figure S6.** Neighbor-joining phylogenetic tree of representative Desulfobulbaceae 16S rRNA gene sequences including the amplified sequence variants (ZOTU) retrieved form Lake Cadagno sediments. ZOTU307 (red circle) was by far the most abundant sequence and exhibited a vertical distribution pattern very similar to that of *Ca.* Methanoperedens. The tree was constructed using Maximum Composite Likelihood correction and partial gap deletion. Bootstrap values >65% for 1000 re-samplings are shown and the scale bar represents 2% estimated sequence divergence.

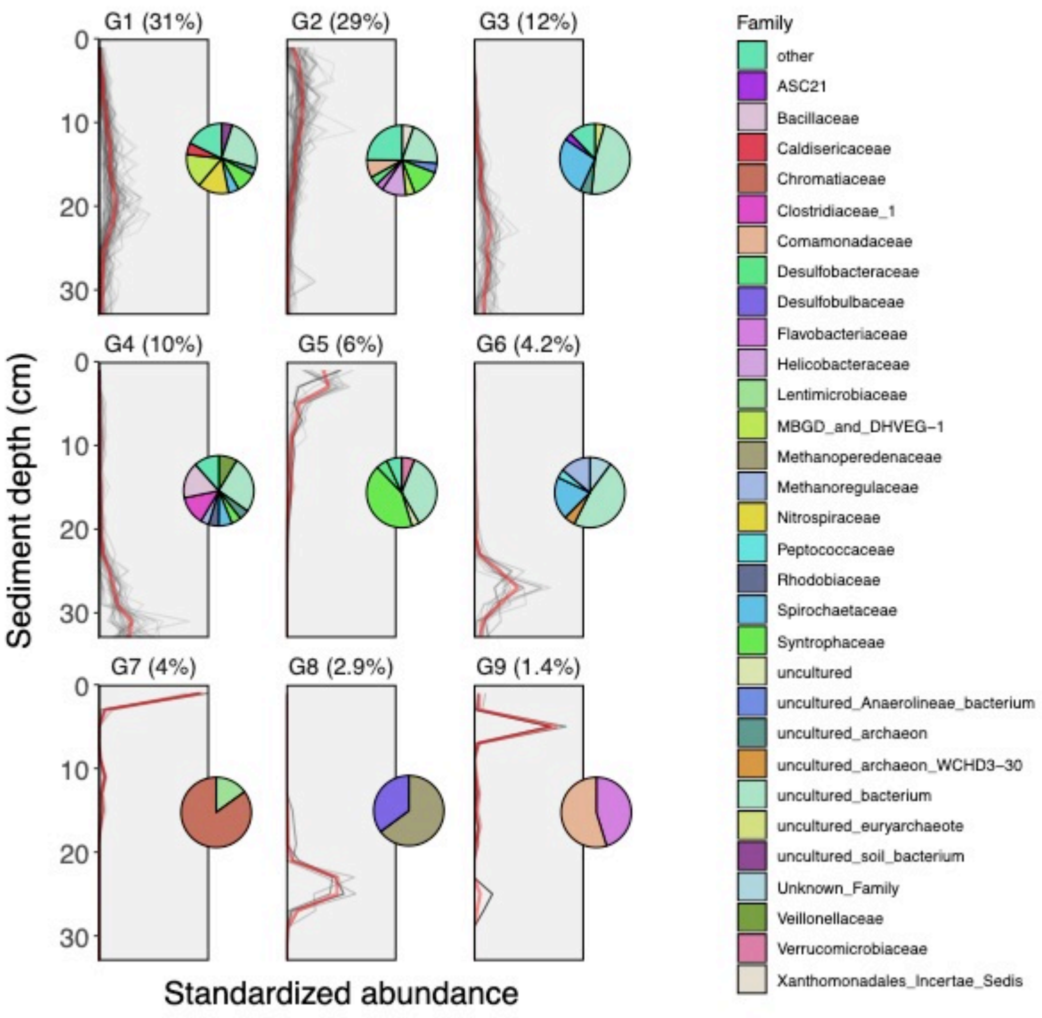


**Figure S7.** In order to test whether other microbial taxa than *Ca.* Methanoperedens and Desulfobulbaceae show a depth distribution pattern matching the subsurface maximum of anaerobic methane oxidation (AOM), we clustered the vertical abundance profiles of 265 ZOTUs, representing 99% of all sequences, into 9 distribution patterns (G1-9). Percentages in parentheses indicate the portion of all sequences falling into a specific grouping. The grey lines represent the profiles of normalized abundances of individual ZOTUs, and the red lines indicates the mean of all ZOTUs in a group. The cluster analysis revealed an AOM-like pattern only for Desulfobaceae, and Methanoperedenaceae (G8). Clustering was done using "ward.D", based on Euclidian distances of normalized vertical abundance profiles. Calculations, clustering and graphs were done using the "plot_groups" function in the R library "phylo.lipids" (Weber et al. 2018) (https://github.com/yukiweber/phylo.lipids).

### Supplementary Tables

Table S1. Components of medium used for $^{13}CH_4$ incubation experiments. All medium components were autoclaved separately and mixed aseptically after cooling. The medium was sulfate-free and modified concentrations of basic salts were used (Ettwig et al. 2009).

| Components | In 10 L medium | Final concentration (mM) |
|---|---|---|
| $KH_2PO_4$ | 0.68 g | 0.5 |
| $MgCl_2.6H_2O$ | 1.015 g | 0.5 |
| $CaCl_2.2H_2O$ | 0.735 g | 0.5 |
| $NaHCO_3$ (20 mM) | 100 ml | 2.0 |
| acid trace [a] | 5 ml | - |
| alkaline trace [b] | 2 ml | - |

[a] 1L 100 mM HCl containing: 2.085 g $FeSO_4·7H_2O$, 0.068 g $ZnSO_4·7H_2O$, 0.12 g $CoCl_2·6H_2O$, 0.5 g $MnCl_2·4H_2O$, 0.32 g $CuSO_4$, 0.095 g $NiCl_2·6H_2O$, 0.014 g $H_3BO_4$

[b] 1L 10 mM NaOH containing: 0.067g $SeO_2$, 0.050g $Na_2WO_4·2H_2O$, 0.242g $Na_2MoO_4$

Table S2. Coverage of the primer pair 515F-Y / 926R for ANME sequences in the SILVA database (SSU-128, RefNr) assessed by the TestPrime tool (Klindworth et al. 2013) (https://www.arb-silva.de/search/testprime/).

| Taxa | | Coverage (%) | Accessions | Match | Mismatch |
|---|---|---|---|---|---|
| Phylum | Family/Genus | | | | |
| Euryarchaeota | ANME-1a | 85.4 | 43 | 35 | 6 |
| Euryarchaeota | ANME-1b | 91.3 | 104 | 94 | 9 |
| Euryarchaeota | ANME-2a/b | 91.2 | 126 | 114 | 11 |
| Euryarchaeota | ANME-2b | 91.7 | 12 | 11 | 1 |
| Euryarchaeota | ANME-2c | 90.6 | 86 | 77 | 8 |
| Euryarchaeota | ANME-3 | 88.9 | 63 | 56 | 7 |
| Euryarchaeota | Methanoperedenaceae (ANME-2d) | 96.2 | 106 | 102 | 4 |

### References


Ettwig, K. F., T. Van Alen, K. T. Van De Pas-Schoonen, M. S. M. Jetten, and M. Strous. 2009. Enrichment and molecular detection of denitrifying methanotrophic bacteria of the NC10 phylum. Appl Env. Microbiol **75**: 3656–3662. doi:10.1128/AEM.00067-09

Klindworth, A., E. Pruesse, T. Schweer, J. Peplies, C. Quast, M. Horn, and F. O. Glöckner. 2013. Evaluation of general 16S ribosomal RNA gene PCR primers for classical and next-generation sequencing-based diversity studies. Nucleic Acids Res. **41**: 1–11. doi:10.1093/nar/gks808

Kostka, J., and K. H. Nealson. 1998. Isolation, cultivation and characterization of iron and manganese-reducing bacteria. Tech. Microb. Ecol.

Kumar, S., G. Stecher, and K. Tamura. 2016. MEGA7: Molecular evolutionary genetics analysis version 7.0 for bigger datasets. Mol. Biol. Evol. **33**: 1870–1874. doi:10.1093/molbev/msw054

Schwertmann, U., and R. M. Cornell. 2000. Iron oxides in the laboratory: Preparation and characterization, Wiley-VCH Verlag GmbH.

Weber, Y., J. S. Sinninghe Damsté, J. Zopfi, and others. 2018. Redox-dependent niche differentiation provides evidence for multiple bacterial sources of glycerol tetraether lipids in lakes. Proc. Natl. Acad. Sci. **115**: 10926–10931. doi:10.1073/pnas.1805186115
